# Spatial processing of limbs reveals the center-periphery bias in high level visual cortex follows a nonlinear topography

**DOI:** 10.1101/2023.10.15.561711

**Authors:** Edan Daniel Hertz, Jewelia K. Yao, Sidney Gregorek, Patricia M. Hoyos, Jesse Gomez

## Abstract

Human visual cortex contains regions selectively involved in perceiving and recognizing ecologically important visual stimuli such as people and places. Located in the ventral temporal lobe, these regions are organized consistently relative to cortical folding, a phenomenon thought to be inherited from how centrally or peripherally these stimuli are viewed with the retina. While this eccentricity theory of visual cortex has been one of the best descriptions of its functional organization, whether or not it accurately describes visual processing in all category-selective regions is not yet clear. Through a combination of behavioral and functional MRI measurements, we demonstrate that a limb-selective region neighboring well-studied face-selective regions defies predictions from the eccentricity theory of cortical organization. We demonstrate that the spatial computations performed by the limb-selective region are consistent with visual experience, and in doing so, make the novel observation that there may in fact be two eccentricity gradients, forming a parabolic topography across visual cortex. These data expand the current theory of cortical organization to provide a unifying principle that explains the broad functional features of many visual regions, showing that viewing experience interacts with innate wiring principles to drive the location of cortical specialization.

## Introduction

High-level visual cortex (HLVC), which spans the ventral and lateral surfaces of the human temporal lobe, comprises clusters of neurons involved in the perception and recognition of ecologically relevant stimuli such as faces, words, places, and limbs^1–4^. These category-selective regions are thought to emerge in an experience-dependent manner across childhood^5–11^. The functional organization of these category-selective regions is strikingly consistent across individuals, whereby each region’s location is anchored to a particular cortical fold^12–14^. In addition to being responsive to a preferred visual stimulus, each region receives information from preferred locations in the visual field^15–18^. This region of visual space from which a neuron responds to a stimulus is called the neuron’s receptive field (RF)^19^. Because visual cortex is organized retinotopically, neighboring neurons have neighboring RFs. In HLVC, neurons located more laterally within ventral temporal cortex (VTC) near the Fusiform gyrus have central receptive fields responding to stimuli near the point of fixation, while those in the medial VTC have receptive fields which sample the periphery of the visual field more heavily^20^. As such, there is a smooth lateral-to-medial eccentricity gradient thought to emerge from simple feed-forward wiring principles and is thus inherited from retinotopic visual field maps in early visual cortex. This interesting and surprisingly consistent pattern gave rise to the prominent Eccentricity Theory for functional organization of HLVC: objects viewed with central vision (e.g., faces) will lead to specialized clusters in lateral VTC (e.g., the lateral Fusiform gyrus), and more eccentric/peripheral ones in medial VTC^6,21^.

Faces, for example, are best recognized with central vision^22,23^. Across development, face-selective regions causally involved in face perception^24^ emerge on the lateral posterior and middle Fusiform gyrus (pFus and mFus) in HLVC (Fig. 1a). Scenes often extend well into our peripheral vision, and the scene-selective region involved in navigation is concordantly located more medially within the collateral sulcus (CoS, Fig. 1a). Interestingly, despite the fact that it abuts face-selective cortex, receptive field properties of a region involved in the perception and recognition of limbs have yet to be mapped. Indeed, the lateral location of the limb-selective region, nestled laterally within the occipitotemporal sulcus (OTS) near face-selective regions, would predict foveally-biased receptive fields (Fig. 1a). However, an examination of how limbs are behaviorally experienced as a visual stimulus might predict the opposite.

**Figure 1:**
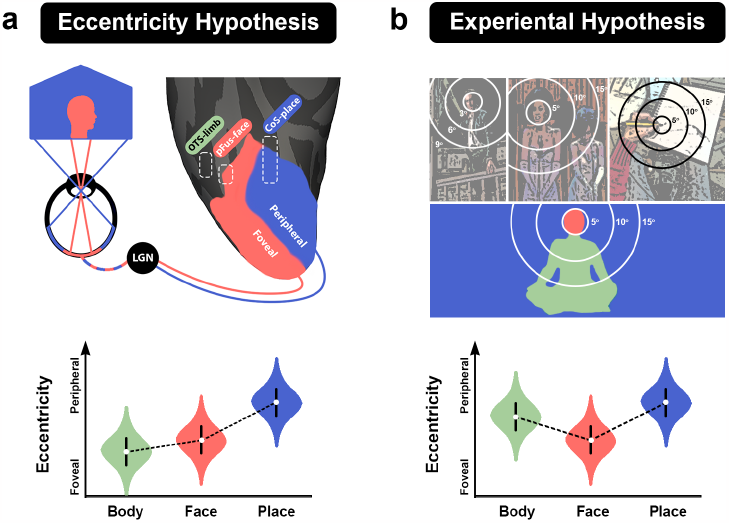
Eccentricity gradient in visual cortex and viewing experience of body and limbs make competing predictions for body-selective receptive field properties. **(a)** The current Eccentricity Hypothesis describes how pRFs located in lateral VTC sample the central visual field, while those located medially are more peripheral. The outline labeled pFus corresponds to the approximate location of a face-selective region on the posterior Fusiform. The CoS outline is a place-selective region in the collateral sulcus. The occipitotemporal sulcus (OTS) outline denotes a body-selective region whose lateral location predicts pRFs with a more foveal bias than pFus, as illustrated with hypothetical distributions of pRF eccentricity in violin plots in the lower inset. **(b)** Viewing behavior by humans in social contexts is heavily biased towards fixating on the face (upper left images), positioning bodies and limbs more peripherally on the retina. Even when interacting with objects (upper right image), one’s own limbs are more peripherally located in the lower visual field. This pattern would predict that pRF properties would pool information from peripheral locations of the lower visual field, as illustrated with hypothetical distributions of pRF eccentricity in violin plots in the lower inset, each category colored according to the inset image of a sitting person.

In the vast majority of social interactions, humans fixate their gaze on the face^25,26^, placing body and limb information more peripherally within the visual field. Even when alone, one’s own limbs during work or ambulation are usually located below the point of one’s fixation. Thus, an experiential account of limbs would predict peripherally located receptive fields within limb-selective cortex (Fig. 1b), a prediction in strong contrast to the long-standing eccentricity theory. Should limb-selective pRFs demonstrate foveal biases, then our account for how visual experience maps onto cortical representation during development would be insufficient. However, peripherally-located pRFs would potentially overturn the current working model of visual cortex organization. These conflicting predictions make the limb-selective region the perfect testing ground for differentiating the eccentricity theory and the experiential account of visual cortex organization.

Differentiating these conflicting theories would require mapping both visual category representation and RF properties within the same individuals. However, most previous studies using checkerboard stimuli for pRF mapping employed high-contrast checkerboard stimuli, a stimulus that is relatively insufficient for driving strong responses in HLVC where limb-selective cortex is located. Extending our recent effort to design high-level retinotopic mapping stimuli^27^, here we created a retinotopic mapping paradigm with rich social content capable of driving visual responses well beyond primary visual cortex to overcome previous hurdles in mapping limb-selective cortex. Combined with naturalistic eye tracking data, we ask: Are pRF properties of limb-selective cortex dictated by their position in the retinal eccentricity gradient (Fig. 1a)? Or, do the spatial computations performed by limb-selective cortex align with the retinal location of limbs as they are experienced visually (Fig, 1b)? In answering these questions, we hope to provide an alternative framework of cortical organization that accounts for the disparate predictions made by eccentricity and experiential accounts of visual cortex.

## Results

### What are the retinal statistics of limbs as they are visually experienced?

A wide body of research posits that visual experiences in childhood have influential effects on cortical development and functional organization in adulthood^5,8,11,28^. In fact, the eccentricity theory posits that biases in the way different stimuli are viewed during development are what leads to the distinct but consistent organization of category-selective regions in HLVC^6,16,20,21,29^. Faces are predominantly viewed with central vision across development^16^, yet how limbs and bodies are visually experienced during naturalistic viewing is less clear. While bodies have been shown to be important for person recognition at a distance^30^, the statistical location of limbs and bodies on the retina compared to other categories like faces or scenes has not been quantified. Given the prominence of faces in social interactions, and the tendency to fixate on faces, we hypothesize that limbs and bodies will consistently result in more peripheral retinal images, particularly in the lower visual field (below fixation on faces).

To test this hypothesis, a total of 25 individuals (average age 23±8.6y, 52% female) participated in a naturalistic eye-tracking study. Each individual viewed 25 videoclips, each 30-seconds long, taken from different Hollywood films^31^. Movie clips were chosen such that they included people (with visible faces and bodies) either alone or engaging in social interactions, with other objects and a background scene present (Fig. 2a). Participants were instructed to watch the clips as they normally would when enjoying a movie with the exception that head motion was controlled through a chin-rest, and fixations were eye-tracked for the duration of the experiment. To assess the retinal image that bodies and limbs produce, a subset of frames were extracted from each movie clip, sampled at 0.2Hz (n=6 frames per clip, total of n=150 frames). Example fixations on selected frames are illustrated in Figure 2a. Each extracted frame was hand-labeled to produce a binarized mask denoting the location of faces and bodies present within each frame (Fig. 2b). Using these binary masks, we simulated the average retinal image during viewing by shifting each frame so that the point of fixation became the new image’s center (Methods). We averaged the fixation-centered images across participants for each frame, such that regions with brighter colors indicate more consistency across participants (Fig. 2c).

**Figure 2:**
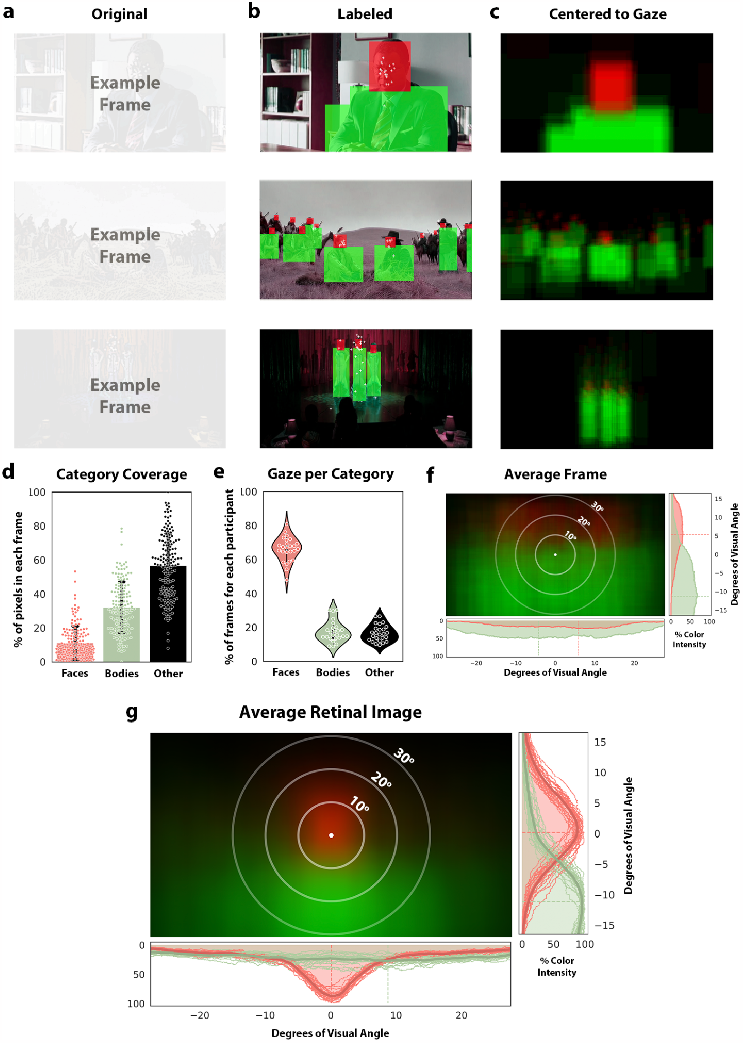
Quantification of viewing experience suggests bodies are mostly viewed in the lower periphery of the visual field. **(a)** Example frames from video clips included in the “Hollywood” dataset^31^, with eye fixations of N=25 participants overlaid as white plus signs. **(b)** Frames were manually labeled with “face” and “body” labels. **(c)** Average “retinal image” from a given frame translated to center the point of fixation, averaged across participants. **(d)** Quantification of how much of the visual field was occupied by faces (red), bodies (blue), or other objects and scenes (black) from extracted movie frames. Each dot is a frame, with a total of 150 frames. **(e)** Violin plots demonstrating the distribution of viewing statistics across all participants and frames. **(f)** The average image across extracted frames, without aligning to fixation. Bodies (blue) statistically occupy the center of the video screen, and faces (red) the upper visual field. **(g)** The average retinal image across all extracted frames and all participants centered according to participant fixation. Increasing intensity of red or blue signifies increased probability of a face or body, respectively, populating that point on the retina. Faces statistically occupy the center of the retina, bodies occupy the lower peripheral visual field.

To quantify these visual statistics, we performed two analyses with matching controls. First, we quantified what proportion of looking time was spent on each visual category by extracting identical moments (i.e., frames) of each movie (n=6 frames per clip). For each participant, we measured the fraction of extracted frames on which a face, a body/limb, or background/object was being fixated (Fig. 2e). We find that faces are the locus of fixation in 66% of the 150 frames (on average over participants), whereas bodies and background/object (‘other’) were fixated on in 17.4% and 16.6% of frames, respectively. That is, we find that faces were more than three times more likely to be the focus of participant fixation compared to either bodies (T[26]=19.6, P<0.001, CI=[43.42, 53.66]; Fig. 2e), or to background/objects (T[26]=23.2, P<0.001, CI=[44.99, 53.79]). One possible confound that is important to address is whether this dramatic difference in gaze location could be explained by the coverage of the frames that each category occupies. That is, could it be that faces are shown more frequently or in larger size than bodies, driving the difference in gaze time between those visual categories? In Figure 2d we quantified the percent of pixels in each frame devoted to each visual category. On average, faces occupied 11% of pixels, bodies 32%, and background and objects (‘other’) 57% of the pixels. That is, even though bodies and limbs occupy three times the pixels that faces do on average (T[149]=15.0, P<0.001, CI=[0.18 0.24]; Fig. 2d), faces are still fixated a factor of three times more often (Fig. 2e).

Second, we sought to quantify the average retinal image produced during movie watching. To produce a model of the inherent visual statistics of the movie frames themselves, we averaged all the labeled video frames. In the average image, consistent patterns across frames should result in regions of solid bright colors, and if there are no consistent patterns across frames, the average image should be completely black, or very blurred. In the average image (Fig. 2f), we observe that bodies tend to be shown in the bottom two thirds of the screen, while faces are more often shown on the top third of the screen, with both categories demonstrating a smooth distribution across the extent of the x-axis. Importantly, we see that the innermost circle in the center of the screen (Fig. 2f) is entirely blue, indicating that bodies most consistently occupy the center of the video frame, more than other categories. Thus, a viewer who was biased towards always fixating near the middle of the screen would be foveating most frequently on body stimuli. In the context of these visual statistics from the video stimuli, we set to quantify the average retinal image by taking into account fixation data. Each extracted video frame was padded and translated so that the point of fixation was centered, and these fixation-centered frames were averaged across all movie clips and participants (Fig. 2g). We can see that faces consistently occupy what would be the fovea, indicated by the bright red region surrounding the point of fixation. Bodies, conversely, were consistently observed outside the point of fixation, and did not begin surpassing faces in their retinal image frequency until about 10-degrees below the point of fixation. When quantifying the differences in stimulus intensity along the vertical meridian between the face (red) and body (green) distributions, we find that the average y-coordinate for peak stimulus intensity of faces was significantly higher in the visual field compared to that of bodies (mean y-coordinate of peak intensity for faces = 0.17°±1.06°, bodies = -11.6°±1.47°; paired T[24]=50.78, P<0.001). Quantifying the difference in x-coordinate of peak stimulus intensity along the horizontal meridian we find a smaller difference between face and body stimuli, where the between-subject variability in peak intensity of body stimulus was much higher than in face stimuli (mean x-coordinate of peak intensity for faces = 0.08°±0.66°, bodies = 8.76°±8.42°; paired T[24]=-3.74, P=0.001). It is worth noting this variability is likely driven by the largely even distribution along the horizontal axis, in which even small biases in fixation will result in a largely shifted peak. Had faces and bodies been fixated on equally during movie watching, we would have seen a region surrounding the fovea showing no preference for either category. However, with faces commanding a large majority of the viewers’ visual attention (Fig. 2e), bodies and limbs, given their physical position below the head, statistically fall beneath the point of fixation, matching well the average retinal image shown in Figure 2g. Taken together, these findings suggest that consistently across individuals and visual scenes, faces tend to be fixated on, while limbs tend to lie in the lower periphery of the retinal image.

### Do limb-selective population receptive fields show a foveal or peripheral bias?

To assess whether the pRF centers of limb-selective cortex are best predicted by their position in the eccentricity gradient (Eccentricity Hypothesis) or viewing behavior (Experiential Hypothesis), we recruited n=27 undergraduate student (17 females, aged 18.6±0.75) to undergo functional MRI while completing two perceptual experiments. The first was a visual localizer presenting images of various categories (faces, bodies, words, houses, objects) while participants underwent functional magnetic resonance imaging (Fig. 3a). The localizer was designed to identify voxels that preferentially respond to images of limbs and bodies (Methods). In each participant, we defined a limb-selective region located within the occipitotemporal sulcus (OTS), at a position approximately half of the sulcus’ length (Fig. 3b). We will refer to this location as OTS-limb^32^, sometimes referred to in the literature as the extrastriate or Fusiform body area (EBA or FBA)^33–35^. Voxels were defined as limb-selective if their response to limbs was statistically greater than all other categories at T-values > 3. An example OTS-limb region is illustrated in two participants in Figure 3b. The eccentricity and experiential hypotheses make predictions about limb-selective pRFs relative to the categories of faces and places: limb selectivity is more lateral than that of faces or places, and limbs are viewed more peripherally than faces in particular. Does the location of OTS-limb on the medial-lateral eccentricity axis, or the relatively peripheral viewing pattern of limbs, determine the pRF properties in the category-selective region? To quantify OTS-limb pRF properties relative to its face- and place-selective counterparts, we identified in every participant the two neighboring face-selective regions located just medially on the posterior and middle Fusiform gyrus, which we refer to as pFus-face and mFus-face^32^, sometimes collectively referred to as the Fusiform Face Area (FFA)^36,37^. We also identified the place-selective region (houses > all other stimuli, T-values>3) located more medially in the collateral sulcus (CoS), referred to here as CoS-place, and sometimes in the literature as the parahippocampal place area (PPA)^38,39^. Segmentations for the three visual categories in two example participants are shown in Figure 3b.

**Figure 3.**
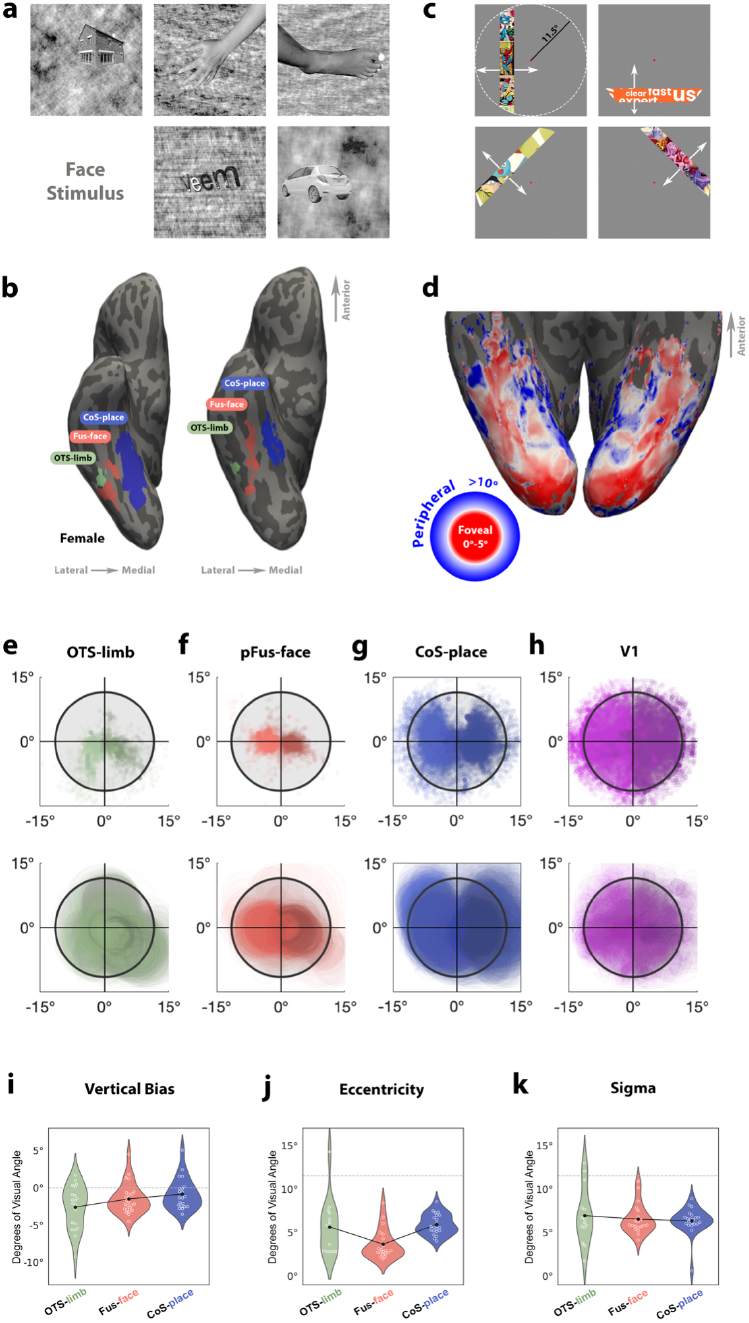
Combined category localization and population receptive field mapping supports an experiential account of high-level visual cortex organization. **(a)** Visual Category Localizer^40^ allowed us to localize category selective regions in each individual brain. Images belonged to six visual categories: houses, hands, feet, faces (adult + child), pseudowords, and cars. **(b)** Right hemisphere of two example subjects in an inferior view, illustrating the labeling of body (green) face (red), and place (blue) selective regions. **(c)** Illustration of the pRF mapping experiment in which a sweeping bar containing flickering cartoon images swept across the screen on the vertical, horizontal, and diagonal axes. Fixation point enlarged for visibility. White arrows indicate directions of motion. **(d)** Cortical map derived from the pRF mapping of a single participant in an inferior view. Each vertex is colored according to the fit pRF’s eccentricity. Warmer colors indicate lower eccentricity (more foveal) while cooler colors represent preference for more eccentric (peripheral) stimuli. Data thresholded by variance explained greater than 10%. **(e-h)** Scatterplots of the pRF centers in each cortical region (top row), and raw pRFs (center+size) from all participants overlaid such that darker areas indicate locations of highest overlap (bottom row). X- and Y- axes show degrees of visual angle. Black circle denotes stimulation presentation area (11.5 degree radius). **(i-k)** Quantification of the pRF properties across regions in individual participants: (i) mean y-ccordinate of pRF centers, (j) mean pRF eccentricity, and (k) mean pRF size, sigma. Violin plots demonstrate distribution of values over participants within each visual category. White circles indicate data from each individual, black dots represent group means, black lines connect group means.

To measure the pRF properties of category-selective voxels, the same participants underwent a retinotopic mapping experiment during fMRI. In a given run, participants fixated on a central dot while a sweeping bar moved across the screen, in which colorful cartoon images flickered at a rate of 7Hz (Fig. 3c), referred to as “Toonotopy”. For each voxel, the population receptive field was modeled as a 2-dimensional gaussian encompassing the area of the visual field that elicited a reliable response from the voxel^41^. The pRF is a stimulus-encoding model, yielding interpretable variables such as the location of the gaussian center (X,Y), its size (σ), and its gain (g). We also fit an exponential term to describe the nonlinearity of the response that occurs in voxels beyond primary visual cortex (V1)^17,42^. In an example participant (Fig. 3d), each vertex of the cortical surface is colored according to the pRF’s radial distance from the point of fixation (e.g., eccentricity). With vertex-wise data thresholded at 10% variance explained, reliable pRFs can be modeled well beyond earlier visual cortex into the anterior temporal lobe. The mean variance-explained across pRFs within category-selective regions is quite high across participants (OTS-limb: 33%±13%; Fus-face: 59%±15%; Cos-place: 50%±11%; V1: 48%±7%). Because the localizer and pRF mapping data are aligned to each individual’s native cortical surface, we can ask how pRF properties differ between category-selective regions.

We find that pRFs in the OTS-limb region (Fig. 3e) are significantly more eccentric and demonstrate a unique upper-lower visual field sampling bias when compared to Fus-face (Fig. 3f). Qualitatively, when pRFs are plotted on the visual field, the OTS-limb region has a denser coverage of the lower visual field compared to Fus-face, whose pRFs are much more centrally biased, huddling around the point of fixation. In contrast, pRFs of the CoS-places region (Fig. 3g) have centers scattered above and below the horizontal meridian, with coverage extending well into the periphery. As a comparison, the receptive fields from V1 are plotted as well, with pRFs more evenly tiling the visual field (Fig. 3h). We quantified differences in receptive field properties (size and eccentricity) of the different category-selective regions, including only vertices that had variance explained of *R*^2^ > 0.1.

First comparing the Y-coordinate of the pRF fits, we find that pRFs of the OTS-limb region have a stronger verticality bias (Fig. 3i), where the average Y-coordinate is located significantly lower in the visual field (2.58 degrees below the point of fixation) compared to Fus-face (1.33 degrees below, T[17]=-2.2, P=0.042, CI=[-2.03,-0.04]) and CoS-place (0.83 degrees below, T[17]=-3.36, P=0.004, CI=[-2.64,-0.60]). We further quantified this bias, asking what fraction of pRFs are located in the lower versus the upper visual field. In each participant we found the percentage of pRFs whose center was in the lower visual field, and compared the percentages in face and limb regions using a paired-samples t-test. While all three category selective regions showed a preference for the lower visual field, we found that in OTS-limbs, the percent of centers in the lower visual field was significantly higher than in CoS-place (Fig. 4d, Limb: 70.09%, Place: 57.31%, T[17]=2.53, P=0.021), and higher than Fus-face, although this difference was not statistically significant (Fig. 4d, Limb: 70.09%, Face: 60.84%, P=0.176).

**Figure 4.**
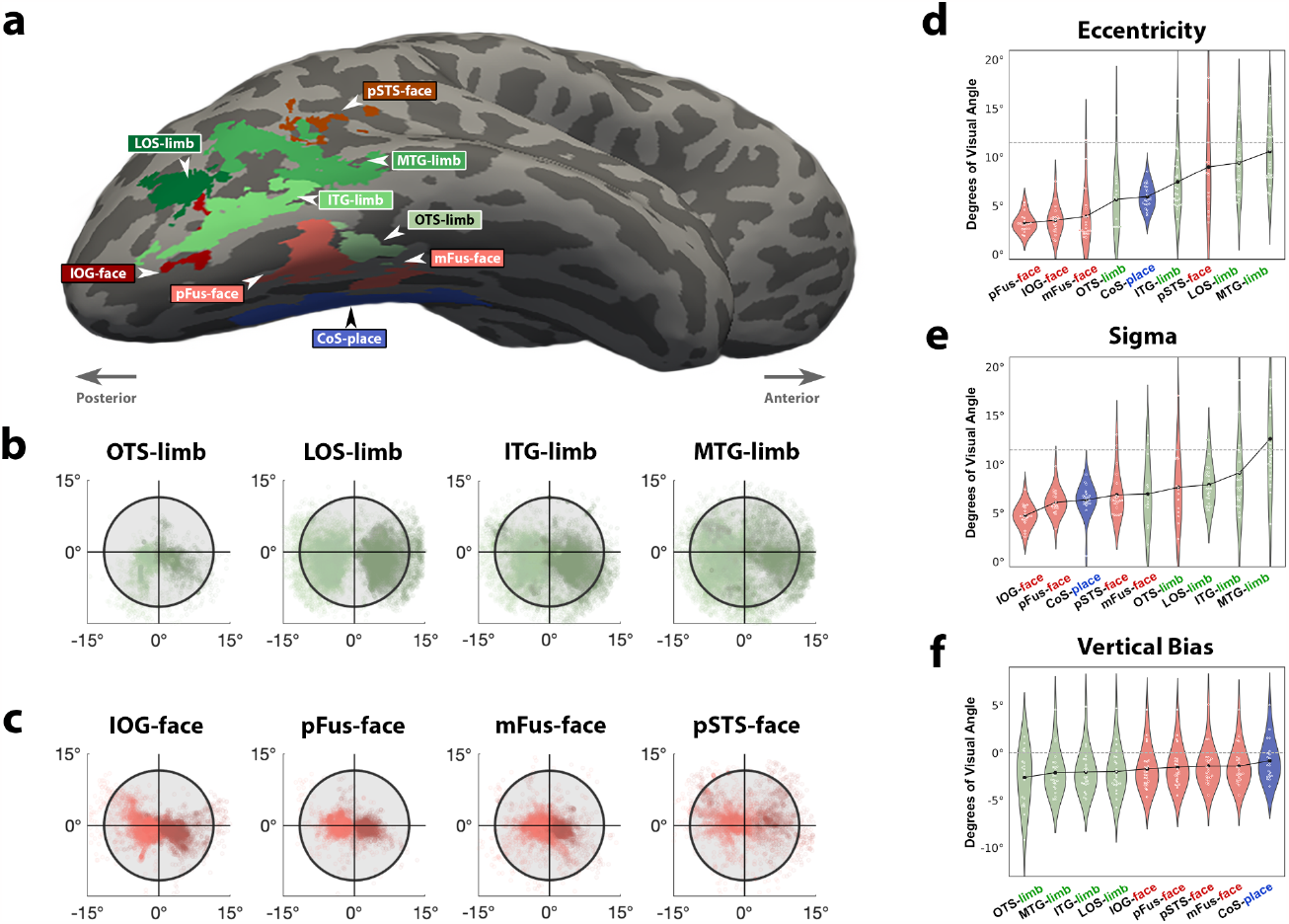
Receptive Field properties of specific ROIs in VTC and the lateral HLVC. (a) Right hemisphere of a single participant showing the spatial distribution of the category selective regions on this participant’s native cortical space. Labels drawn based on results from visual localizer using anatomical landmarks for accurate labeling in native brain space. (b-c) Scatterplots depicting locations of pRF centers in each ROI in the limb (b) and face (c) selective regions. x- and y- axes are in degrees of visual field. Dot: the spatial location of a pRF center from a single vertex from a single participant. Dots are semi-transparent to visualize pRF center overlap. Within each plot, darker colors indicate data from the Left hemisphere, light colors from the Right. Black circle marks stimulus borders at 11.5 degrees. (d-f) Violin plots depicting pRF eccentricity (d), size (e), and vertical bias (f) across ROIs in the HLVC. White circles indicate individual data, black dots mark group mean in each ROI, and black lines connect group means. X-axis order ranked by value on y-axis, showing the regions with lowest average y-value (d), eccentricity (e) or size (f) on the left, and regions with the highest values on the right. (f) Vertical bias in each ROI quantified as the average y-value of pRF centers in each participant.

Comparing pRF properties across visual categories, we found that pRFs in OTS-limb were significantly more eccentric than in the Fus-face region (Fig. 3j, Limb: 5.8 degrees, Face: 3.6 degrees, T[13]=3.36, P=0.005, CI=[1.01,4.89]). We found no significant difference in the size of the pRFs across those visual regions (Limb: 7.01 degrees, Face: 6.49 degrees, T[13]=1.01, P=0.33, CI=[-0.89, 2.44]). In sum, combining pRF mapping with a visual category fMRI localizer, we demonstrate that limb-selective cortex in HLVC contains distinctly peripheral receptive fields biased towards sampling the lower visual field when compared to face-selective regions. These findings are surprisingly inconsistent with the longstanding eccentricity theory, and are more closely aligned with predictions made by the experiential account for functional development of the HLVC.

### Peripheral bias is a general feature of both limb-selectivity as well as lateral temporal cortex

One test of the experiential hypothesis is that a limb-selective region should show a peripheral and lower-visual field bias regardless of its cortical location. In the lateral visual stream^43^, there are three additional limb-selective regions which surround motion-selective cortex (area hMT+) in the ascending portion of the inferior temporal sulcus ^44^. To test if these three regions also show a peripheral and lower visual field bias, we defined in the same participants the additional limb-selective regions ^45^ located on the inferior temporal gyrus (ITG-limb), the lateral occipital sulcus (LOS-limb), and the middle temporal gyrus (MTG-limb). Spatial distribution of those regions is shown in an example participant in Figure 4a. Extracting the pRFs from each region’s voxels using the same pRF-modeling protocol, we can plot the fit receptive field centers on the visual field (Fig. 4b). It is readily visible that all three lateral limb-selective regions have, like the ventral OTS-limb, pRF centers that show a bias towards sampling the lower visual field, but extend even further into the periphery. Extending this analysis to additional lateral face-selective regions on the inferior occipital gyrus (IOG-faces) and the posterior superior temporal sulcus (pSTS-faces) we find that pRF centers on these more lateral face-selective regions are also more peripherally-biased than their ventral face-selective counterparts (Fig. 4c).

Indeed, ranking these category-selective regions by their mean pRF eccentricity reveals that the lateral pSTS-faces is more peripherally-biased than ventral face regions (mFG, pFG), and that all of the lateral limb-selective regions, together with pSTS-faces, are the most peripherally-biased (Fig. 4d). While consistent with previous observations of pSTS-faces being less foveally biased than ventral regions ^27,46^, the observation that pSTS-faces more heavily samples the visual periphery, when viewed in the context of the peripheral bias of lateral limb-selective regions, suggests that peripheral visual field coverage may be a general feature of lateral visual cortex organization. In fact, this effect is so extreme that all the lateral limb-selective regions and pSTS-faces show a larger bias towards the peripheral visual field than even the CoS-place region (Fig. 4d), a region previously hallmarked by its peripheral visual bias. When ranked by mean pRF size, we see a categorical split: ventral face-selective regions cluster together with the smallest pRF sizes compared to limb-selective regions showing the largest (Fig. 4e). Ranking category-selective regions by their verticality bias (mean Y-coordinate of pRF centers) reveals an additional categorical split, regions cluster together with the largest preference for the lower visual field in limb-selective regions, strongest in the OTS-limb region, followed by a moderate lower bias in face-selective regions, and lastly CoS-place with the least vertical bias (Fig. 4f).

### A single eccentricity gradient is not sufficient to explain category-selective pRF properties

The peripheral bias of the OTS-limb region defies the foveal prediction made by the eccentricity hypothesis, demonstrating that the current working hypothesis mapping visual experience to cortical representation is insufficient. However, the same data offer a parsimonious solution for extending the eccentricity hypothesis to account both for the OTS-limb region’s seemingly paradoxical cortical location and the phenomenon of peripheral bias in lateral category-selective regions. From the perspective of the lateral Fusiform gyrus, receptive fields increase in size and eccentricity when one travels not only medially along the cortical surface, but also laterally. Does this suggest that there are two eccentricity gradients emanating from the lateral Fusiform gyrus (FG)- one medially, and one laterally?

To test this hypothesis in an observer-independent fashion, we parcellated visual cortex into seven anatomically-defined ROIs according to cortical folds, with the medial-most ROI defined by the collateral sulcus and the lateral-most ROI defined by the middle temporal gyrus (Fig. 5a). Anatomical ROIs were drawn in a common cortical space (i.e., fsaverage), and all participants’ eccentricity maps were transformed into this space using cortex-based alignment. We thresholded voxels for variance-explained values exceeding R^2^ > 0.1, and calculated the mean eccentricity of pRFs within each anatomical ROI, for each participant. When the eccentricity values of each ROI are plotted in order along the lateral to medial axis, a clear parabolic gradient emerges with pRFs in the most lateral and medial ROIs demonstrating higher eccentricity values than those located near the middle, around the Fusiform gyrus (Fig. 5b). A repeated measures (within-subject) ANOVA indicated that eccentricity values differ significantly across anatomical ROIs, in both hemispheres (Right: F(6)=18.3, P<0.001; Left: F(6)=26.46, P<0.001). To better understand the pattern of change in eccentricity values along the medial-lateral axis, we performed a model comparison analysis, fitting the data with both a linear model and a quadratic model, and comparing the goodness-of-fit between those models. We found that the fit of the quadratic model was better than that of the linear model with higher variance-explained and lower root mean square error in both hemispheres (Right-linear: *R*^2^ = 0.19, *RMSE*= 2. 96, Right-quadratic: *R*^2^ = 0. 24, *RMSE* = 2. 87; Left-linear: *R*^2^ = 0. 24, *RMSE* = 3. 85, Left-quadratic: *R*^2^ = 0. 34, *RMSE* = 3.6), providing additional, quantitative support for a parabolic eccentricity pattern, compared to a linear gradient of eccentricity. These data offer strong evidence that the eccentricity gradient previously described in the literature extending from the collateral sulcus to the lateral Fusiform gyrus reverses when one continues traveling laterally to mid-temporal regions on the lateral cortical surface.

**Figure 5:**
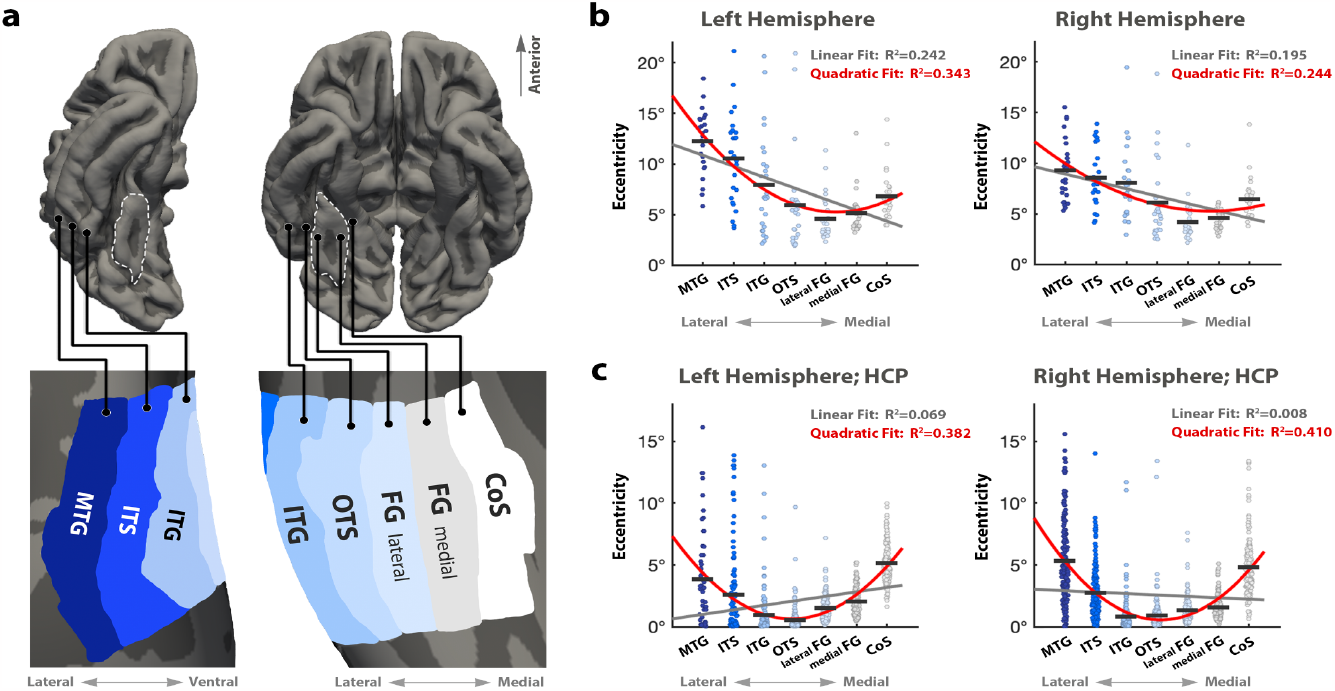
Parabolic Eccentricity Gradient in Across High Level Visual Cortex (HLVC). **(a)** Labels in fsaverage space separate the HLVC into ROIs based on anatomical landmarks along the medial-lateral axis. Top row shows pial surface view, bottom row zooms into labeled regions on an inflated surface for clarity. **(b)** Swarm plots depicting eccentricity values across the medial-lateral axis from 27 participants. Dots represent average eccentricity from a single participant in a single ROI. Black horizontal lines represent mean eccentricity per ROI across participants. Gray line shows fit of linear model, red line shows fit of quadratic model. **(c)** Swarm plots depicting the same analysis performed on the HCP 7T dataset (n=181). Labels, from medial to lateral: CoS- Collateral Sulcus; FG medial- Medial Fusiform Gyrus; FG lateral- Lateral Fusiform Gyrus; OTS- Occipital-temporal sulcus; ITG- Inferior temporal gyrus; ITS- Inferior temporal sulcus; MTG- Mid temporal gyrus.

To test the robustness of this finding, we used data from the HCP Retinotopy dataset ^47^ which employed different stimulus-mapping techniques in addition to higher field-strength fMRI. We repeated our analysis to test if our results would replicate in this larger (n=181), independent sample. Because the HCP data is in fsaverage space, we could use the same anatomical labels to repeat our previous analysis on this independent dataset. Using the same pipeline, we find strikingly similar and significant results: the eccentricity values are lowest (most foveal) between the OTS and FG and become more eccentric with increasing distance from the FG either medially or laterally. We ran a within-subject, repeated measures ANOVA and found that in both hemispheres, eccentricity values differ significantly across anatomical ROIs (Right: F(6)=107.8, P<0.001; Left: F(6)=21.96, P<0.001). In the model comparison analysis we found that the goodness of fit of the quadratic model was much higher than that of the linear model in both hemispheres (Right-linear: *R*^2^ = 0. 008, *RMSE* = 2. 63, Right-quadratic: *R*^2^ = 0. 41, *RMSE* = 2. 03; Left-linear: *R*^2^ = 0. 07, *RMSE* = 2. 4, Left-quadratic: *R*^2^ = 0. 382, *RMSE* = 1. 96). Taken together, these findings suggest that a parabolic pattern of eccentricity is a general, replicable organizational principle across ventral and lateral visual streams.

## Discussion

The functional organization of visual category-selective regions is strikingly consistent across individuals, whereby each region’s location is anchored to a particular cortical fold ^12,13^. One prominent theory aiming to explain the principle that governs this consistent organization of high-level visual cortex (HLVC) is the Eccentricity Theory, which posits that objects viewed with central vision will lead to specialized clusters on the lateral VTC, and more eccentric/peripheral ones on the medial VTC ^6,21^, which show preferential connectivity to foveal and peripheral representation of primary visual cortex. Previous studies supporting the Eccentricity Theory have mainly used an experimental design which effectively stimulates VTC, but is insufficient for driving reliable responses in HLVC to properly fit pRF models. Thus, the lateral HLVC encompassing the limb-selective region has not yet been characterized using pRF mapping techniques. In this study, we demonstrate that pRFs in a limb-selective region are drastically more peripheral than the foveal predication made by the Eccentricity Theory. Using eye-tracking, we were able to quantify the statistics of visual experiences, showing that limbs are most commonly processed using peripheral vision. Our findings suggest that our visual experience drives the biases in spatial computations of high-level vision. Seeking to produce a parsimonious model of visual cortex organization, we quantified the graded changes in pRF eccentricity more broadly beyond the ventral surface, from the medial collateral sulcus (CoS) to the lateral mid-temporal gyrus (MTG). We found that the cortex near the lateral Fusiform gyrus (FG) is a point of transition where the eccentricity gradient reverses, forming a parabolic eccentricity gradient, with foveal representations in the FG and peripheral representations in the most medial and lateral regions of high-level visual cortex. This primary description of mapped receptive fields supports a novel organization principle of the high level visual cortex in which the ventral and lateral streams of visual processing may not be as separate as once described, but instead linked by a shared and parabolic gradient of receptive field properties.

The first evidence for a cortical region that responds selectively to hands was observed in single cell recordings in macaque IT cortex ^3,48^. This groundbreaking finding suggesting that cells in IT cortex are sensitive to such high-level categories was not immediately accepted, with the first replication of the finding published a full decade later ^49^. In the decades to follow, the scientific community produced countless studies focusing on face-selective regions in both animal models and human fMRI studies, while limb/body-selective regions were left understudied. In the early 2000s, fMRI studies of the responses to body stimuli lead to the novel definition of the Extrastriate-Body Area (EBA), a large region in the lateral occipitotemporal cortex (LOTC), located near the motion-sensitive MT cortex ^35^. Follow up fMRI studies have revealed that limb/body selectivity is not exclusive to the LOTC, with additional regions identified near the the mid Fusiform gyrus (mFG; ^50,51^, as well as in the posterior superior temporal sulcus (pSTS; ^52^, later being parcellated more finely according to their anatomical locations ^45,53^. Despite sharing the same consistent relationship to cortical folding as face- and place-selective regions, limb-selective regions were never formally included in models describing this consistent relationship between category selectively and cortical folding, such as the Eccentricity Theory.

While this theory makes a simple prediction for limb-selective cortex–a more lateral position in the OTS should predict more foveally-biased pRFs compared to face regions–this spatial bias would be at odds with how bodies and limbs tend to be viewed during social interactions. Previous work measuring neural responses demonstrated that social stimuli including faces and bodies elicit the largest responses when presented in their typically-experienced orientations^54^. Coupled with observations showing that faces are powerful attractors of attention and eye gaze^25,55^, and make up a dense proportion of central visual input within the first few months of life^56^ all suggest that bodies and limbs, as visual stimuli, largely enter the visual system through the periphery of the retinal image. The peripheral retinal sampling of limbs and bodies in social contexts match well the empirical data from pRF model fits produced across two datasets in the present study. That is, limb-selective pRFs were significantly more eccentric than the centrally-biased fits of face-selective cortex. Had there truly been a single eccentricity gradient dictating the location of a given category-selective region based on the viewing statistics of that category during development^6,20,21^, then limb-selective cortex should appear more medially in VTC around the medial Fusiform gyrus. Its lateral location thus suggests visual experience plays a strong role in determining the receptive field properties of limb selectivity, joining a wide body of research positing that sensory experiences in childhood have influential effects on cortical development and functional organization in adulthood ^5–10^. This is not to say that the OTS-limb region does not sample foveal information at all, indeed some of its receptive fields do overlap with the fovea which is consistent with work showing that the body is an important visual source for recognition when the face is not available^30^. Our observations suggest that the more peripheral input experience during social interactions and video watching make up a larger percentage of visual input. Thus, while the position of OTS-limb defies the original Eccentricity Theory’s prediction for its receptive field properties, we offer here a parsimonious extension of the theory to describe both ventral and lateral aspects of high-level visual cortex simultaneously. Rather than a single, linear gradient of eccentricity extending medially from the collateral sulcus and ending at the lateral Fusiform gyrus, we show here evidence that this gradient demonstrates a reversal in pattern around the lateral Fusiform, with pRF eccentricity increasing again as one traverses the cortical mantle towards the lateral surface, ending at the middle temporal gyrus. This parabolic eccentricity gradient better describes this larger swatch of cortex, and like some previous models describing object size and animacy preferences^57^, is capable of comprehensively capturing visual selectivity patterns for any arbitrary piece of occipitotemporal cortex.

Despite being one of the last category-selective regions to have its RF properties described, the limb-selective region of the OTS is influential given that it was at odds with current theories of visual cortex organization. The current eccentricity theory describing the organization of category-selective regions in the ventral temporal lobe was rooted in the notion that feed-forward axonal wiring principles from early retinotopic maps creates an input bias, with lateral regions receiving foveal input and medial ones-peripheral input. By extending recent observations that a face-selective region on the lateral temporal surface shows strong peripheral connectivity^27,46^, we proposed an extension of the eccentricity theory whereby cortical regions beyond the lateral Fusiform show a reversal for preferred retinotopic input and sample from increasingly peripheral locations of the retinal image, ultimately forming a parabolic gradient between the collateral sulcus of the medial ventral temporal lobe and the middle temporal gyrus of the lateral temporal lobe. The idea that there are two increasing eccentricity gradients sharing a common foveal border is consistent with previous observations in visual cortex that visual field maps share a border at a reversal of polar angle gradients^58,59^. While visual field map borders are defined by polar angle reversals, the existence of multiple, discrete foveal representations shared by clusters of visual field maps in early and high-level visual cortex^58^ necessarily implies that eccentricity representations should also show such reversals, albeit along a perpendicular dimension. Indeed, our observed parabolic eccentricity gradient runs medial-lateral, perpendicular to the direction of phase-map reversals of the ventral visual stream^60^. In this case, the reversal of an eccentricity gradient, which seems to happen over larger swaths of cortex, might represent broader-scale functional transitions, such as separation points between the ventral, lateral, and dorsal streams of visual processing^43,61–63^.

While this broader parabolic organization is likely inherited from axonal wiring set up during gestation^64^, it does not have to be the only feature that determines the cortical location and properties of a category-selective region, given that these category regions receive not only axonal input from earlier visual cortex (e.g., via inferior longitudinal pathways^65^) but receive and contribute axons to the vertical occipital^66,67^ (VOF) and the Arcuate^68^ (AF) fasciculi. Indeed, the area around the OTS appears to be a transition zone: despite limb-selective cortex, word-selective cortex, and potentially other learned categories like Pokémon all cohabitating this cortical fold, words^15^ and Pokémon^6^ show foveally-biased pRF sampling. Given that the perceptual goal of reading words or recognizing learned game characters may be distinct from the perceptual goal of processing limbs, it is likely that signals relayed through other major white matter pathways will play a major role in the spatial properties of developing pRFs. For example, word-selective cortex, and thus its receptive fields, likely depend on language lateralization^69,70^ rather than purely visual processes alone. Thus, while the parabolic eccentricity gradient we describe seems to be a broad cortical feature that plays a role in the gross functional organization in high-level vision, it is likely not the only feature contributing to pRF properties. Attention, for example, is one top-down factor that plays a role in scaling visual responses in ventral visual cortex^71^, and can shift the spatial properties of receptive fields^17,72^. In the current dataset, participants were instructed to fixate on the center but attend to the sweeping bar, whereas in the HCP dataset^73^ they were attending centrally. This attentional difference may partially explain some differences between the datasets, but nonetheless demonstrates that the parabolic gradient we present here is robust to changes in attention. Future research should study how these additional top-down signals originating from parietal and frontal cortices further refine and bias pRF properties across development.

This study offers the first characterization of the receptive field properties (RF) in limb-selective regions of high-level visual cortex, demonstrating that those RFs are lower, more peripheral, and generally larger than RFs in face-selective regions. Using a novel pRF mapping experiment we were able to map limb-selective RF properties in lateral aspects of HLVC which have yet to be characterized, establishing the groundwork for laying out a new hypothesis for functional organization of category selective regions across visual cortex more broadly, including its most lateral aspects. Our findings suggest that the eccentricity gradient from medial to lateral VTC flips and extends towards the lateral HLVC in a parabolic fashion, with most foveal receptive fields located near the lateral Fusiform gyrus with decreasing central bias in two directions: medially, and laterally.

## Methods

### Participants

A total of 27 participants aged 18-20 (mean±sd 18.6±0.75y, 17 females) participated in the fMRI study. One additional participant could not remain awake during fMRI, and was thus excluded from analyses. A total of 25 separate participants (age mean±sd 23±8.6y, 13 females) participated in the eye-tracking experiment. Two additional participants were excluded from the eye-tracking dataset due to insufficient data collected due to technical difficulties. All participants were right-handed with normal or corrected-to-normal vision, and provided informed, written consent to participate in the experiment. Procedures were approved by the Princeton Internal Review Board on human subjects research. Retinotopy data from the publicly available HCP dataset were collected from a total of 181 subjects (109 females), as described in the original publication ^74^.

#### Data Acquisition

MRI and fMRI data were collected at the Scully Center for the Neuroscience of Mind and Behavior within the Princeton Neuroscience Institute, using a Siemens 3-Tesla Skyra system. Each participant completed a single recording session in which they completed four runs of a Visual Category Localizer Experiment ^40^ (4m each), followed by a qMRI anatomical scan ^75,76^ (25m), a break outside the scanner (5m), and three runs of the pRF Mapping Experiment (5m each).

##### Quantitative magnetic resonance imaging (qMRI) acquisition

An artificial T1-weighted image was produced using the following qMRI protocol for the purposes of reconstructing the cortical surface. Quantitative MRI was used to estimate T1 relaxation times based on existing protocols ^75,76^, measured from three spoiled gradient echo (spoiled-GE) images with flip angles of 4°, 10°, and 20°. An artificial T1-weighted anatomical 3D image was then produced from these measurements for each participant for the purposes of segmenting white and gray matter through FreeSurfer software.

##### Functional MRI

For the visual category localizer, fMRI scans comprised 72 slices acquired using a multiplexed echo planar imaging (EPI) sequence (multiband acceleration factor: 3, voxel size: 2.5mm isotropic), with repetition time (TR) = 2s, echo time (TE) = 3000ms, and flip angle (FA) = 80°. Each run was 209s in duration and there were four runs in total. For the retinotopic mapping experiment, scan parameters were identical with the exception of multiband acceleration factor set to 2 and 48 total slices.

##### Eyetracking data

The behavioral experiment involving movie-watching was conducted within the Scully Center for the Neuroscience of Mind and Behavior using an SR Research EyeLink 1000 Plus system. Visual stimuli were displayed on an Asus LCD monitor (PG278QR: 2560×1440p, 120Hz refresh rate, ultra-low motion blur enabled), and controlled with an Apple Mac Pro running OSX 10.11. The stimulus monitor measures 597mm wide, located 570mm from the viewer’s eyes, resulting in a retinal image of 55 by 33 degrees of visual angle. Participants’ eyes were calibrated using 5 point calibration (four corners, one center) and validated before movie viewing began. Eye gaze position was sampled and recorded at a rate of 1000Hz.

#### Eyetracking Hollywood Movies Experiment

Participants (n=25) viewed 25 videoclips of 30-second duration, taken from different Hollywood films^31^. Movie clips were chosen such that they included people (with visible faces and bodies) either alone or engaging in social interactions, with other objects and a background scene present (Fig. 2a) to allow participants the option to fixate on a range of animate and inanimate stimuli in any given clip. Participants were instructed to watch movies as they normally would when enjoying a movie. Participant head motion was controlled using a table-mounted chin-rest. To efficiently analyze data, a subset of frames were extracted from each movie clip at a rate of 0.2Hz, resulting in 6 frames samples per clip, for a total of 150 frames. The same frames were extracted across participants to enable the quantification of the consistency of the retinal image produced during a given scene. Face and body stimuli were hand-labeled using rectangles that subtended the vertical and horizontal extent of the pixels occupied by the face or body, where faces were assigned RGB values of [255,0,0] and bodies were assigned RGB values of [0,255,0]. All other pixels were defined as black [0,0,0] such that each pixel of each frame was categorized as face (red), body (green), or background/other (black). To produce a simulation of the retinal image within a participant according to their point of fixation at a given frame, the extracted frame was padded with RGB values of zeros, the image was centered/translated such that the point of fixation was brought into the center of the frame, and then the image was cropped back to the dimensions of the viewing monitor.

#### Visual Category Localizer

During each run (209s) participants fixated on a central dot as visual stimuli were presented in 4s blocks. Each block consisted of eight images alternating at 2Hz, from a single visual category out of six in total: houses, hands, feet, faces (adult + child), pseudowords, and cars (Fig. 1a). Limb stimuli included both upper and lower limbs, and always included the digits. Each image was tightly cropped and overlaid on a textured grayscale background produced by phase scrambling a stimulus from another category, where an additional baseline condition consisted of a scrambled background with no overlaid image ^40^. Images subtended a visual angle of 7.125° centered on the fovea and were presented with the Psychophysics Toolbox ^77^ using code written in MATLAB (The Mathworks, Natick, MA). To ensure attention to the stimuli, participants completed a 2-back task, where they were instructed to press a button whenever they detected that an image repeated itself with exactly one intervening image between repeats.

#### Population Receptive Field Mapping

During each run (300s), participants fixated on a central dot as a bar (2° visual angle) swept across the screen on the vertical, horizontal, and diagonal axes with two directions of motion, revealing portions of a large circle defining the full visual field (radius = 11.5°; 8 bar sweeping configurations; Fig. 3c). The bar displayed a strip of colorful images from a selection of >100 colorful cartoons and comics, alternating at a rate of 7Hz. This experiment is referred to as “Toonotopy”. At regular intervals, the apertures were removed and participants viewed a mean luminance gray background with a central fixation point. The cartoon stimuli were chosen to represent a wide array of stimuli including bodies, faces, words, houses and objects. This experiment was adapted from our previous work^27^, with changes to the stimulus content (more evenly sampling different categories), image refresh rate (7Hz rather than 8Hz), and bar motion (smooth, continuous sweeping motion rather than discrete 2-degree steps). Another important difference from the previous version of Toonotopy is that participants attended to the visual stimulus (e.g., the sweeping bar) and monitored it for the appearance of wiggling bumble bees. Sweeping bars appeared on a low-contrast, radial grid resembling a spider web and participants were instructed to detect the bees to help Charlotte the Spider remove them from her web. This narrative was designed specifically for a separate pediatric experiment, but was also given to the present adult participants for consistency.

#### Anatomical Data Analysis

QMRI data were processed using the mrQ software package in MATLAB to produce T1-weighted maps ^76^. The full analysis pipeline and its published description is freely accessible at (https://github.com/mezera/mrQ). The QMRI-derived artificial T1-weighted anatomical image was produced for each individual participant, segmented with FreeSurfer (https://surfer.nmr.mgh.harvard.edu/), and then passed through iBeat for an optimal separation of gray and white matter^78,79^ and fed back to FreeSurfer to correct the white matter segmentation. The resulting segmentations were used for cortical surface reconstruction^80,81^ and visualization of retinotopic data in each individual’s native brain space.

#### Functional Data Analysis

Analyses were performed in each individual’s native brain space. An inflated cortical surface reconstruction was used for each participant and all functional data (localizer and pRF data) were projected onto this cortical surface in which voxel time-series data were resampled to each cortical vertex. Data was preprocessed based on the HCP minimal preprocessing pipeline for motion-correction, slice-timing correction, and lastly phase-distortion correction using Topup^82,83^. No spatial smoothing was used. Resampling voxel time-series data to cortical vertices and general linear model (GLM) analyses were processed through FsFast (FreeSurfer Functional Analysis Stream; https://surfer.nmr.mgh.harvard.edu/) to derive for each voxel the beta-weights associated with each category for the production of contrast maps, described below.

#### Labeling of category selective regions

Category-selective regions of interest (ROIs) were defined for each individual participant using data from the visual category localizer experiment. For labeling each category selective region, statistical contrasts of the category of interest > all other stimuli were thresholded at T-values greater than 3 and overlaid onto the cortical surface for each subject. The borders around each of the category selective regions were drawn on the cortical surface, and the vertices within the border above a threshold of 3 were defined as that region’s ROI for that participant.

For limb-selective regions, we used the statistical contrast of [hands > non-limb stimuli]. The borders for each limb-selective ROI were determined using anatomical landmarks as detailed by ^45^, and included limb-selective regions on the occipitotemporal sulcus (OTS-limb), lateral occipital sulcus (LOS-limb), inferotemporal gyrus (ITG-limb), and middle temporal gyrus (MTG-limb). For localization of face-selective regions, we used the contrast [face > all other stimuli], and included face-selective regions on the inferior occipital gyrus (IOG-face), the posterior Fusiform (pFus-face), the middle Fusiform gyrus (mFus-face), and the posterior branch of the superior temporal sulcus (pSTS-face). For localization of the place-selective region, we used the contrast [house > all other stimuli] and included a single place-selective region on the collateral sulcus (CoS-place). All labeled regions are shown on an example right hemisphere of a single participant in Figure 4a.

#### Population receptive field (pRF) Modeling

Using the fMRI data acquired from the sweeping bar task, we performed population receptive field (pRF) mapping ^41,42,84–87^, which is a non-invasive, computational modeling technique that allows quantification of pRF properties in each voxel. Sweeping the stimulus across the visual field while the participant maintains central fixation allows us to estimate what subset of the visual field each voxel is most sensitive to. A receptive field is modeled for every voxel independently as a 2D gaussian with a center (**x**,**y**) and a size determined by σ, the standard deviation term in the gaussian model. Through an iterative process optimizing the gaussian properties, the predicted and observed fMRI time-series are compared to find the pRF parameters that best predict the observed time-series for that voxel by minimizing the root-mean-square-error. Software repository for pRF modeling was developed by the Stanford VISTA Lab (github.com/vistalab) ^41^, with additions for compressive nonlinearity implemented by Kendrick Kay ^42^, as well as adaptations by the NYU Winawer Lab for use in FreeSurfer-style data formats (github.com/WinawerLab/prfVista)^88^.

In regards to visualizing pRF data, receptive field centers and sizes can be visualized for each participant in each region of interest (ROI) based on the modeled pRFs using custom made plots. Coverage plots are commonly used to compare the average pRF between ROIs, but averaging pRFs can result in a spurious coverage that does not reflect the actual properties of the underlying pRFs. Thus, aiming to minimize the use of mass averaging methods while presenting raw data whenever possible, in Figure 3e (bottom row) we plot an overlay of all the raw pRFs (x,y,σ) from all subjects on a single plot, such that darker regions indicate regions of significant overlap. The limitation of this visualization is that individuals that have more data points have more influence on the resulting image as the number of modeled vertices across participants is not constant, but no participant can contribute more than 500 vertices in this approach to minimize visualization bias. All statistical quantifications were random-effects analyses in which pRF properties from a given individual were averaged before entering statistical models. Finally, in Figure 3e (top row) we plot only the centers of pRFs to emphasize the spread of the pRFs across the visual field in the different visual regions.

#### Statistics

Receptive field data were fit using a stimulus-encoding model^41^ to derive parameters describing the circular gaussian model fit for each vertex on the cortical surface. No spatial smoothing was performed on functional MRI data. In the fMRI analysis, the data used to define the ROIs were independent of the data used to calculate the pRF properties (visual category localizer task and pRF mapping task, respectively), to avoid circularity confounds. When comparing pRF parameters between ROIs, within-subjects statistics were used, such as paired t-tests and within-subjects ANVOAs. Statistical analyses comparing pRFs were done using random-effects models, with each participant contributing a single data point summarizing a given pRF property from a cortical region. Where possible, all individual data points were shown directly (e.g., swarm plots) or shown within visual summary illustrations (e.g., violin plots). Data visualizations were made using custom-built scripts in MATLAB (The Mathworks, Natick, MA), as well as using Matplotlib^89^ and Seaborn^90^ Python packages. Resulting p-values quantifying statistical significance were always two-tailed. The degrees of freedom for each analysis (or number of participants within an analysis) are reported following the relevant statistic within the text. Data were assumed to be normal, with no gross violations of normality observed. Given that data points are compared in a within-subjects manner with no grouping of participants, group randomization was not relevant in this particular study. No statistical methods were used to predetermine sample sizes but our sample sizes are similar to those reported in previous publications using HCP data, as well as in previous work^91^.

## References

1. Downing, P. E., Chan, A. W.-Y., Peelen, M. V., Dodds, C. M. & Kanwisher, N. Domain specificity in visual cortex. Cereb. Cortex 16, 1453–1461 (2006).

2. Cohen, L. et al. The visual word form area. Brain vol. 123 291–307 Preprint at 10.1093/brain/123.2.291 (2000).

3. Gross, C. G., Rocha-Miranda, C. E. & Bender, D. B. Visual properties of neurons in inferotemporal cortex of the Macaque. J. Neurophysiol. 35, 96–111 (1972).

4. Pinsk, M. A., DeSimone, K., Moore, T., Gross, C. G. & Kastner, S. Representations of faces and body parts in macaque temporal cortex: a functional MRI study. Proc. Natl. Acad. Sci. U. S. A. 102, 6996–7001 (2005).

5. Arcaro, M. J., Schade, P. F., Vincent, J. L., Ponce, C. R. & Livingstone, M. S. Seeing faces is necessary for face-domain formation. Nat. Neurosci. 20, 1404–1412 (2017).

6. Gomez, J., Barnett, M. & Grill-Spector, K. Extensive childhood experience with Pokémon suggests eccentricity drives organization of visual cortex. Nat Hum Behav 3, 611–624 (2019).

7. Thompson, A., Gribizis, A., Chen, C. & Crair, M. C. Activity-dependent development of visual receptive fields. Curr. Opin. Neurobiol. 42, 136–143 (2017).

8. Golarai, G., Liberman, A. & Grill-Spector, K. Experience Shapes the Development of Neural Substrates of Face Processing in Human Ventral Temporal Cortex. Cereb. Cortex 27, bhv314 (2015).

9. Huber, E., Mezer, A. & Yeatman, J. D. Neurobiological underpinnings of rapid white matter plasticity during intensive reading instruction. Neuroimage 243, 118453 (2021).

10. McKyton, A., Ben-Zion, I., Doron, R. & Zohary, E. The Limits of Shape Recognition following Late Emergence from Blindness. Curr. Biol. 25, 2373–2378 (2015).

11. Scherf, K. S., Behrmann, M., Humphreys, K. & Luna, B. Visual category-selectivity for faces, places and objects emerges along different developmental trajectories. Dev. Sci. 10, F15–30 (2007).

12. Weiner, K. S. et al. The mid-fusiform sulcus: a landmark identifying both cytoarchitectonic and functional divisions of human ventral temporal cortex. Neuroimage 84, 453–465 (2014).

13. Grill Spector, K., Weiner, K. S., Kay, K. N. & Gomez, J. The Functional Neuroanatomy of Human Face Perception. Annual Review of Vision Science 3, (2017).

14. Natu, V. S. et al. Sulcal Depth in the Medial Ventral Temporal Cortex Predicts the Location of a Place-Selective Region in Macaques, Children, and Adults. Cereb. Cortex 31, 48–61 (2021).

15. Le, R., Witthoft, N., Ben-Shachar, M. & Wandell, B. The field of view available to the ventral occipito-temporal reading circuitry. J. Vis. 17, 6 (2017).

16. Gomez, J., Natu, V., Jeska, B., Barnett, M. & Grill-Spector, K. Development differentially sculpts receptive fields across early and high-level human visual cortex. Nat. Commun. 9, 788 (2018).

17. Kay, K. N., Weiner, K. S. & Grill-Spector, K. Attention reduces spatial uncertainty in human ventral temporal cortex. Curr. Biol. 25, 595–600 (2015).

18. Silson, E. H., Chan, A. W., Reynolds, R. C., Kravitz, D. J. & Baker, C. I. A Retinotopic Basis for the Division of High-Level Scene Processing between Lateral and Ventral Human Occipitotemporal Cortex. J. Neurosci. 35, 11921–11935 (2015).

19. Hubel, D. H. & Wiesel, T. N. Receptive fields and functional architecture of monkey striate cortex. J. Physiol. 195, 215–243 (1968).

20. Levy, I., Hasson, U., Avidan, G., Hendler, T. & Malach, R. Center–periphery organization of human object areas. Nat. Neurosci. 4, 533–539 (2001).

21. Hasson, U., Levy, I., Behrmann, M., Hendler, T. & Malach, R. Eccentricity bias as an organizing principle for human high-order object areas. Neuron 34, 479–490 (2002).

22. Poltoratski, S., Kay, K., Finzi, D. & Grill-Spector, K. Holistic face recognition is an emergent phenomenon of spatial processing in face-selective regions. Nat. Commun. 12, 4745 (2021).

23. Hsiao, J. H.-W. & Cottrell, G. Two fixations suffice in face recognition. Psychol. Sci. 19, 998–1006 (2008).

24. Rangarajan, V. & Parvizi, J. Functional asymmetry between the left and right human fusiform gyrus explored through electrical brain stimulation. Neuropsychologia 83, 29–36 (2016).

25. Yarbus, A. L. Eye Movements and Vision. (Springer, 2013).

26. Amso, D., Haas, S. & Markant, J. An eye tracking investigation of developmental change in bottom-up attention orienting to faces in cluttered natural scenes. PLoS One 9, e85701 (2014).

27. Finzi, D. et al. Differential spatial computations in ventral and lateral face-selective regions are scaffolded by structural connections. Nat. Commun. 12, 2278 (2021).

28. Srihasam, K., Vincent, J. L. & Livingstone, M. S. Novel domain formation reveals proto-architecture in inferotemporal cortex. Nat. Neurosci. 17, 1776–1783 (2014).

29. Malach, R., Levy, I. & Hasson, U. The topography of high-order human object areas. Trends in Cognitive Sciences vol. 6 176–184 Preprint at 10.1016/s1364-6613(02)01870-3 (2002).

30. Rice, A., Phillips, P. J., Natu, V., An, X. & O’Toole, A. J. Unaware person recognition from the body when face identification fails. Psychol. Sci. 24, 2235–2243 (2013).

31. Costela, F. M. & Woods, R. L. A free database of eye movements watching ‘Hollywood’ videoclips. Data in Brief 25, 103991 (2019).

32. Weiner, K. S. & Grill-Spector, K. Neural representations of faces and limbs neighbor in human high-level visual cortex: evidence for a new organization principle. Psychol. Res. 77, 74–97 (2013).

33. Taylor, J. C., Wiggett, A. J. & Downing, P. E. Functional MRI analysis of body and body part representations in the extrastriate and fusiform body areas. J. Neurophysiol. 98, 1626–1633 (2007).

34. Peelen, M. V. & Downing, P. E. The neural basis of visual body perception. Nat. Rev. Neurosci. 8, 636–648 (2007).

35. Downing, P. E., Jiang, Y., Shuman, M. & Kanwisher, N. A cortical area selective for visual processing of the human body. Science 293, 2470–2473 (2001).

36. Puce, A., Allison, T., Gore, J. C. & McCarthy, G. Face-sensitive regions in human extrastriate cortex studied by functional MRI. J. Neurophysiol. 74, 1192–1199 (1995).

37. Kanwisher, N., McDermott, J. & Chun, M. M. The fusiform face area: a module in human extrastriate cortex specialized for face perception. J. Neurosci. 17, 4302–4311 (1997).

38. Epstein, R. & Kanwisher, N. A cortical representation of the local visual environment. Nature 392, 598–601 (1998).

39. Aguirre, G. K., Zarahn, E. & D’Esposito, M. An area within human ventral cortex sensitive to ‘building’ stimuli: evidence and implications. Neuron 21, 373–383 (1998).

40. Stigliani, A., Weiner, K. S. & Grill-Spector, K. Temporal Processing Capacity in High-Level Visual Cortex Is Domain Specific. J. Neurosci. 35, 12412–12424 (2015).

41. Dumoulin, S. O. & Wandell, B. A. Population receptive field estimates in human visual cortex. Neuroimage 39, 647–660 (2008).

42. Kay, K. N., Winawer, J., Mezer, A. & Wandell, B. A. Compressive spatial summation in human visual cortex. J. Neurophysiol. 110, 481–494 (2013).

43. Pitcher, D. & Ungerleider, L. G. Evidence for a Third Visual Pathway Specialized for Social Perception. Trends Cogn. Sci. 25, 100–110 (2021).

44. Dumoulin, S. O. et al. A new anatomical landmark for reliable identification of human area V5/MT: a quantitative analysis of sulcal patterning. Cereb. Cortex 10, 454–463 (2000).

45. Weiner, K. S. & Grill-Spector, K. Not one extrastriate body area: Using anatomical landmarks, hMT+, and visual field maps to parcellate limb-selective activations in human lateral occipitotemporal cortex. Neuroimage 56, 2183–2199 (2011).

46. Pitcher, D. et al. The Human Posterior Superior Temporal Sulcus Samples Visual Space Differently From Other Face-Selective Regions. Cereb. Cortex 30, 778–785 (2020).

47. Benson, N. C. et al. The Human Connectome Project 7 Tesla retinotopy dataset: Description and population receptive field analysis. J. Vis. 18, 23 (2018).

48. Gross, C. G., Bender, D. B. & Rocha-Miranda, C. E. Visual Receptive Fields of Neurons in Inferotemporal Cortex of the Monkey. Science vol. 166 1303–1306 Preprint at 10.1126/science.166.3910.1303 (1969).

49. Richmond, B. J., Wurtz, R. H. & Sato, T. Visual responses of inferior temporal neurons in awake rhesus monkey. J. Neurophysiol. 50, 1415–1432 (1983).

50. Peelen, M. V. & Downing, P. E. Selectivity for the human body in the fusiform gyrus. J. Neurophysiol. 93, 603–608 (2005).

51. Schwarzlose, R. F., Baker, C. I. & Kanwisher, N. Separate face and body selectivity on the fusiform gyrus. J. Neurosci. 25, 11055–11059 (2005).

52. Pinsk, M. A. et al. Neural representations of faces and body parts in macaque and human cortex: a comparative FMRI study. J. Neurophysiol. 101, 2581–2600 (2009).

53. Weiner, K. S., Sayres, R., Vinberg, J. & Grill-Spector, K. fMRI-Adaptation and category selectivity in human ventral temporal cortex: Evidence for the scaling and sharpening models. Journal of Vision vol. 9 466–466 Preprint at 10.1167/9.8.466 (2010).

54. Chan, A. W., Kravitz, D. J., Truong, S., Arizpe, J. & Baker, C. I. Cortical representations of bodies and faces are strongest in commonly experienced configurations. Nat. Neurosci. 13, 417–418 (2010).

55. Cerf, M., Frady, E. P. & Koch, C. Faces and text attract gaze independent of the task: Experimental data and computer model. J. Vis. 9, 10.1–15 (2009).

56. Jayaraman, S., Fausey, C. M. & Smith, L. B. Why are faces denser in the visual experiences of younger than older infants? Dev. Psychol. 53, 38–49 (2017).

57. Konkle, T. & Caramazza, A. Tripartite organization of the ventral stream by animacy and object size. J. Neurosci. 33, 10235–10242 (2013).

58. Wandell, B. A., Dumoulin, S. O. & Brewer, A. A. Visual field maps in human cortex. Neuron 56, 366–383 (2007).

59. Sereno, M. I. et al. Borders of multiple visual areas in humans revealed by functional magnetic resonance imaging. Science 268, 889–893 (1995).

60. Wang, L., Mruczek, R. E. B., Arcaro, M. J. & Kastner, S. Probabilistic maps of visual topography in human cortex. Cereb. Cortex 25, 3911–3931 (2015).

61. Goodale, M. A. & Milner, A. D. Separate visual pathways for perception and action. Trends Neurosci. 15, 20–25 (1992).

62. Weiner, K. S. & Gomez, J. Third Visual Pathway, Anatomy, and Cognition across Species. Trends Cogn. Sci. 25, 548–549 (2021).

63. Ungerleider, L. G. & Haxby, J. V. ‘What’ and ‘where’ in the human brain. Curr. Opin. Neurobiol. 4, 157–165 (1994).

64. Kamps, F. S., Hendrix, C. L., Brennan, P. A. & Dilks, D. D. Connectivity at the origins of domain specificity in the cortical face and place networks. Proc. Natl. Acad. Sci. U. S. A. 117, 6163–6169 (2020).

65. Tusa, R. J. & Ungerleider, L. G. The inferior longitudinal fasciculus: a reexamination in humans and monkeys. Ann. Neurol. 18, 583–591 (1985).

66. Takemura, H. et al. A Major Human White Matter Pathway Between Dorsal and Ventral Visual Cortex. Cereb. Cortex 26, 2205–2214 (2016).

67. Yeatman, J. D. et al. The vertical occipital fasciculus: a century of controversy resolved by in vivo measurements. Proc. Natl. Acad. Sci. U. S. A. 111, E5214–23 (2014).

68. Catani, M. & Thiebaut de Schotten, M. A diffusion tensor imaging tractography atlas for virtual in vivo dissections. Cortex 44, 1105–1132 (2008).

69. Cai, Q., Paulignan, Y., Brysbaert, M., Ibarrola, D. & Nazir, T. A. The left ventral occipito-temporal response to words depends on language lateralization but not on visual familiarity. Cereb. Cortex 20, 1153–1163 (2010).

70. Dundas, E. M., Plaut, D. C. & Behrmann, M. The joint development of hemispheric lateralization for words and faces. J. Exp. Psychol. Gen. 142, 348–358 (2013).

71. Bugatus, L., Weiner, K. S. & Grill-Spector, K. Task differentially modulates the spatial extent of category-selective regions across anatomical locations. J. Vis. 15, 1170 (2015).

72. Sprague, T. C. & Serences, J. T. Attention modulates spatial priority maps in the human occipital, parietal and frontal cortices. Nat. Neurosci. 16, 1879–1887 (2013).

73. Benson, N. C. & Winawer, J. Bayesian analysis of retinotopic maps. Elife 7, (2018).

74. Benson, N., Jamison, K., Vu, A., Winawer, J. & Kay, K. The HCP 7T Retinotopy Dataset: A new resource for investigating the organization of human visual cortex. J. Vis. 18, 215–215 (2018).

75. Mezer, A., Rokem, A., Berman, S., Hastie, T. & Wandell, B. A. Evaluating quantitative proton-density-mapping methods. Hum. Brain Mapp. 37, 3623–3635 (2016).

76. Mezer, A. et al. Quantifying the local tissue volume and composition in individual brains with magnetic resonance imaging. Nat. Med. 19, 1667–1672 (2013).

77. Brainard, D. H. The Psychophysics Toolbox. Spat. Vis. 10, 433–436 (1997).

78. Li, G. et al. Measuring the dynamic longitudinal cortex development in infants by reconstruction of temporally consistent cortical surfaces. Neuroimage 90, 266–279 (2014).

79. Li, G. et al. Construction of 4D high-definition cortical surface atlases of infants: Methods and applications. Med. Image Anal. 25, 22–36 (2015).

80. Dale, A. M., Fischl, B. & Sereno, M. I. Cortical surface-based analysis. I. Segmentation and surface reconstruction. Neuroimage 9, 179–194 (1999).

81. Fischl, B. FreeSurfer. Neuroimage 62, 774–781 (2012).

82. Andersson, J. L. R., Skare, S. & Ashburner, J. How to correct susceptibility distortions in spin-echo echo-planar images: application to diffusion tensor imaging. Neuroimage 20, 870–888 (2003).

83. Smith, S. M. et al. Advances in functional and structural MR image analysis and implementation as FSL. Neuroimage 23 Suppl 1, S208–19 (2004).

84. Wandell, Winawer & Kay. Computational modeling of responses in human visual cortex. Brain mapping 651–659 (2015).

85. Wandell, B. A. & Winawer, J. Computational neuroimaging and population receptive fields. Trends Cogn. Sci. 19, 349–357 (2015).

86. Tootell, R. B. et al. Functional analysis of V3A and related areas in human visual cortex. J. Neurosci. 17, 7060–7078 (1997).

87. Smith, A. T., Singh, K. D., Williams, A. L. & Greenlee, M. W. Estimating receptive field size from fMRI data in human striate and extrastriate visual cortex. Cereb. Cortex 11, 1182–1190 (2001).

88. Himmelberg, M. M. et al. Comparing retinotopic maps of children and adults reveals a late-stage change in how V1 samples the visual field. Nat. Commun. 14, 1561 (2023).

89. Hunter. Matplotlib: A 2D Graphics Environment. 9, 90–95 (2007).

90. Waskom, M. seaborn: statistical data visualization. J. Open Source Softw. 6, 3021 (2021).

91. Gomez, J. et al. Development of population receptive fields in the lateral visual stream improves spatial coding amid stable structural-functional coupling. Neuroimage 188, 59–69 (2019).

